# Homosalate boosts the release of tumor-derived Extracellular Vesicles with anti-anoikis properties

**DOI:** 10.1101/2021.10.25.465564

**Authors:** Eleonora Grisard, Aurianne Lescure, Nathalie Nevo, Maxime Corbé, Mabel Jouve, Gregory Lavieu, Alain Joliot, Elaine Del Nery, Lorena Martin-Jaular, Clotilde Théry

**Author notes:** Co-last authors (Lorena Martin-Jaular, Clotilde Théry).

## Abstract

Eukaryotic cells, including cancer cells, secrete highly heterogeneous populations of extracellular vesicles (EVs). EVs could have different subcellular origin, composition and functional properties, but tools to distinguish between EV subtypes are scarce. Here, we tagged CD63- or CD9-positive EVs secreted by triple negative breast cancer cells with Nanoluciferase enzyme, to set-up a miniaturized method to quantify secretion of these two EV subtypes directly in the supernatant of cells. We performed a cell-based high-content screening to identify clinically-approved drugs able to affect EV secretion. One of the identified hits is Homosalate, an anti-inflammatory drug found in sunscreens which robustly increased EVs’ release. Comparing EVs induced by Homosalate with those induced by Bafilomycin A1, we discovered that: 1) the two drugs act on EVs generated in distinct subcellular compartments and 2) EVs released upon treatment with Homosalate, but not with Bafilomycin A1, conferred anti-anoikis properties to another recipient tumor cell line. In conclusion, we identified a new drug modifying EV release and demonstrated that under influence of different drugs, triple negative breast cancer cells release EV subpopulations from different subcellular origins harboring distinct functional properties.

## Introduction

Extracellular Vesicles (EVs) are membrane enclosed particles secreted by all types of cells. Given that EVs can transport nucleic acids, proteins and lipids, they are fundamental means of inter-cellular communication [1][2]. EVs can originate in different locations within cells: exosomes originally form as intraluminal vesicles (ILVs) of multivesicular bodies (MVBs) along the endocytic pathway, whereas ectosomes (or microparticles) are generated by direct budding away from the plasma membrane (PM). Exosomes have the same diameter range as ILVs, i.e. 50-150nm, but PM-derived ectosomes, other EVs originating from different compartments (e.g. recycling endosomes), and even some particles released by virus-infected or apoptotic cells can also be in the same size range [1][3]. By contrast, only ectosomes, oncosomes and apoptotic bodies can be also larger (up to 5μm in diameter) [4]. Importantly, different EVs present different combinations of protein markers, as revealed by comparative proteomic studies [5][6][7], and consequently could have distinct functional properties [8][9][10]. In particular, EVs released by cancer cells can play opposite roles in cancer progression [8]. For example, EVs released by breast cancer cells have been shown to display pro-metastatic properties [11][12], similar to those released by pancreatic cancer [13], prostate cancer [14] or liver cancer [15]. Conversely, other studies EVs have reported a protective role of EVs against cancer progression [16][17].

Isolating and studying individual EV subtypes is therefore crucial. Yet, current EV isolation techniques achieve only a partial separation of EV subtypes, although combinations of one or more techniques can considerably improve the distinction between EV subpopulations [1]. The tetraspanins CD63 and CD9 are often used independently as exosome markers. However, they have preferential (but not exclusive) subcellular locations: CD63 is mostly localized in MVBs and consequently more enriched in exosomes, whereas CD9 is mostly localized at the PM and thus more enriched in ectosomes [18], although they can also be expressed simultaneously on the same EVs [5]. Using fluorescent tags to label EV markers (e.g. CD63 or CD9 tetraspanins) can help to study separately the biogenesis mechanisms of EVs containing those markers [18][19]. As an alternative to fluorescent tags, also enzymatic tags can be used to label EVs, like luciferase enzymes [20]. In particular, Nanoluciferase (Nluc) has been successfully used to tag CD63 positive EVs [21].

Here, we tagged CD63 or CD9 [18] with the Nluc enzyme to separately study two populations of EVs (CD63-positive and CD9-positive) released by MDA-MB-231 triple negative breast cancer cells. We then set-up a cell-based high-content screening (HCS) assay to detect the secretion of Nluc-CD63 or Nluc-CD9 tagged EVs by quantifying Nluc activity in the supernatant of cells. HCS methods are extensively used to rapidly identify small molecules with potential biological functions, and are of special interest for repurposing drug applications when performed using chemical libraries compounds already approved by regulatory agencies (FDA, EMA …) [22]. Here, by screening an FDA- and EMA-drug library (Prestwick Chemicals V3), we found that Homosalate acts as a booster of EV secretion in MDA-MB-231 and other tumor cell lines.

We found that Homosalate and Bafilomycin A1, which has been shown to increase the secretion of MVB-derived EVs (=exosomes) by basifying MVB internal pH [23][24][25], acted on distinct subcellular compartments. Homosalate specifically increased an EV subpopulation enriched in a combination of SLC3A2/CD98 and CD9 markers. Importantly, we showed that EVs released by MDA-MB-231 cells upon treatment with Homosalate, but not Bafilomycin A1, conferred anoikis resistance to recipient tumor cells. In conclusion, a novel multi-drug HCS assay for EVs’ release allowed the identification of a never described drug able to increase the secretion of a specific EV subpopulation.

## Results

### Nanoluciferase tagged CD63 and CD9 are secreted into EVs

Our first goal was to develop a quantitative HCS assay to enable consistent and easy read-out of EV secretion in miniaturized 96- and 384-cell culture plate format. We selected two of the most commonly used EV markers, CD63 and CD9 fused to the highly sensitive enzyme Nluc to monitor EV release using stably-expressing cells. We generated constructs encoding for human CD63 or CD9 tagged with Nluc at the N-terminal [26], i.e. away from the C-terminal lysosome-targeting signal of CD63 [27] (Fig. 1A). This luciferase enzyme is very small and exceptionally bright [28], representing the ideal tool for the quantification of EV secretion, even from small amounts of cells. In a bulk population of Nluc-CD63-transfected MDA-MB-231 triple negative breast cancer cells, we observed that Nluc activity was detectable from 390 cells and in the extracellular medium (i.e. supernatant) of 1560 cells (Suppl. Fig. 1A). Thus, we generated stable clonal populations of MDA-MB-231 transfected with Nluc-CD63 or Nluc-CD9. For each cell line, we selected a clone with high Nluc activity in cells (clone 7 for Nluc-CD63 and clone 3 for Nluc-CD9: around 10^8^ luciferase activity/ 25000-30000 cells), and reliable detection in supernatant (Suppl. Fig.1B). For both clones, comparable ratios of supernatant vs cells Nluc activity were observed (6%, Suppl. Fig. 1B). Next, we determined the precise contribution of EVs in the total Nluc activity detected in the supernatant of cells. We used an EV-isolation approach based on Size Exclusion Chromatography (SEC) to separate secreted EVs, which do not enter the pores of the gel and get out first of the column [29], from soluble proteins that could be a potential source of Nluc activity in our system. To accurately characterize the content of SEC fractions from Nluc-CD63 and Nluc-CD9 clones, we compared particle number, protein concentration and Nluc activity in all SEC fractions separately from fraction 7 (F7, first fraction post-void volume of the column) to F24. As shown in Figure 1B and 1C, for both cell lines, particle concentration measured by Nanoparticle Tracking Analysis (NTA) peaked in F9-11, although some particles were still detectable until F16-17 (red line in upper panel), indicating that they are present also in later fractions than in the predicted F7-11. Proteins were detectable from F14 on, with a peak in F20-21 (green line in upper panel), confirming a minimal overlap with EV-enriched fractions. The distribution of Nluc-activity measured in the same fractions (Fig. 1B-C, blue line) closely overlapped with particle concentration distribution, with a peak in F9-11 and some detection until F16-17. Interestingly, a smaller peak appeared in F21-22 corresponding to the peak of free proteins and indicating that a tiny portion of Nluc activity source is free in the supernatant or associated to very small membrane-derived objects (Fig.1B-C). According to these observations, we defined “EV-rich” fractions F7-15 and “Protein-rich” fractions F16-24. In this pilot experiment, we calculated the percentage of Nluc activity in EV-rich (F7-15) and Protein-rich (F16-24) over the total Nluc-activity detected in all fractions (F7-24). We determined that 68% and 71% of the total Nluc activity in Nluc-CD63 and Nluc-CD9, respectively, was associated to EV-rich fractions (Fig. 1B-1C). Conversely, protein-rich fractions were the source of Nluc activity for only 32% or 29% of the total Nluc activity in Nluc-CD63 (Fig.1B) and Nluc-CD9 (Fig. 1C). To further confirm EV specific enrichment in the fractions F7-15 we checked by Western Blot the presence of EV markers in all SEC fractions from Nluc-CD63 and Nluc-CD9 clones. As expected, we found that proteins were highly enriched in “Protein-rich” F16-24 and barely detectable in “EV-rich” F7-15 (Fig. 1D-E). Importantly, EV markers SLC3A2/CD98 (http://exocarta.org/index.html) [18], CD63, CD9 and Syntenin were mainly enriched in F8-15 with peaks in F9, F10 and F11, consistent with previous findings (Fig. 1B and 1C). Conversely, 14-3-3, a protein commonly identified in sEVs recovered by ultracentrifugation (http://exocarta.org/index.html) but recently defined as non-exosomal marker [6] was found at low level in F7-15 and instead more enriched in F16-24 (Fig. 1D and 1E), suggesting that 14-3-3 cannot be used as a EV marker. Finally, pools of SEC fractions from Nluc-CD63 or Nluc-CD9 conditioned media were analyzed by Transmission Electron Microscopy (TEM). We confirmed that the majority of CD63- or CD9-positive EVs in Nluc-CD63 and Nluc-CD9, respectively, were detected in F7-11 (Fig. 1F-G), which correspond to the peak of Nluc activity.

**Figure 1.**
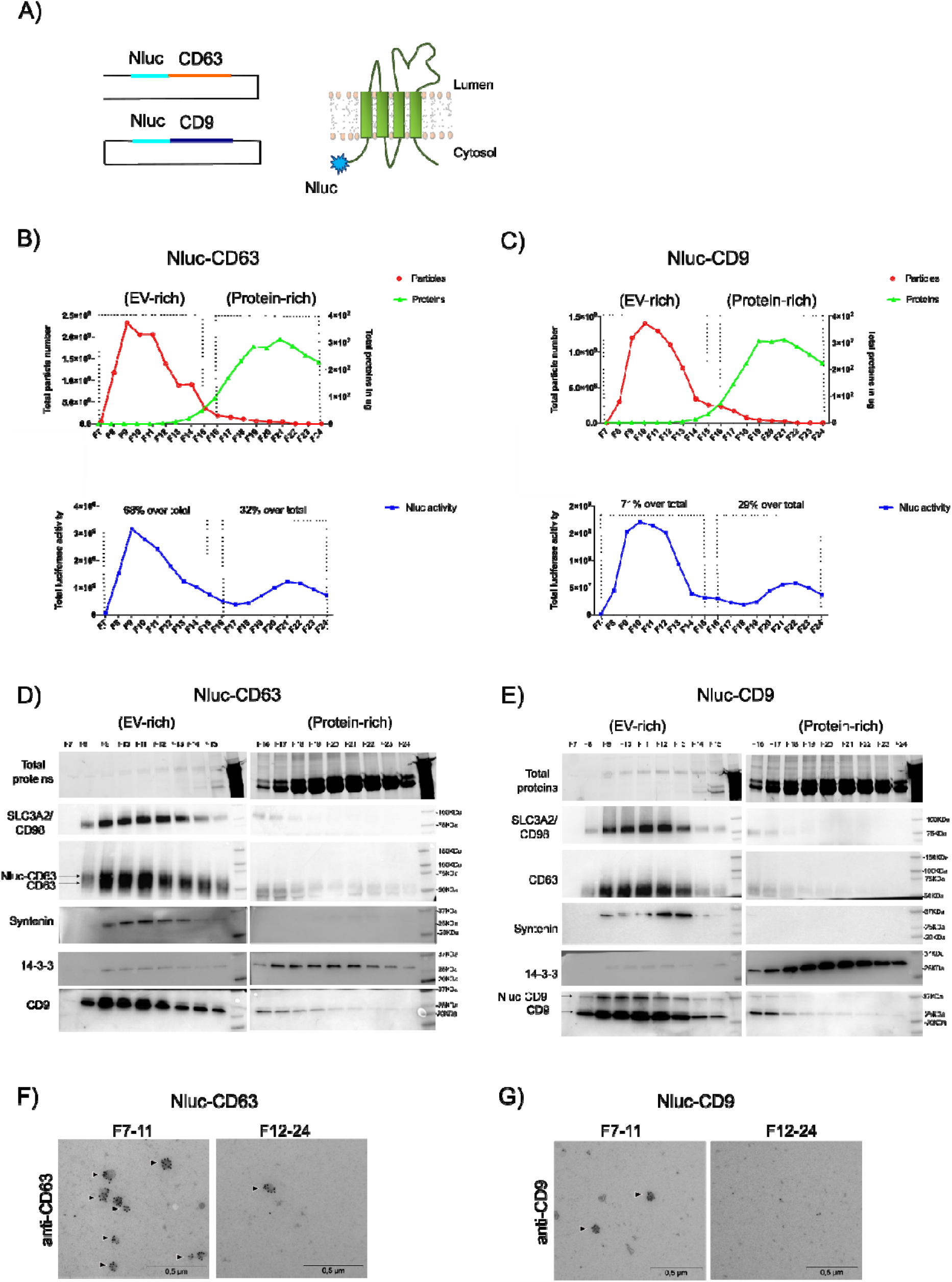
Nanoluciferase tagged CD63 and CD9 are secreted into EVs. A) Left: scheme of Nluc-CD63 and Nluc-CD9 plasmid constructs. Nluc enzyme was cloned into CD63 or CD9 encoding plasmids at the N-terminal position. Right: scheme of the topology of a tetraspanin (CD63 or CD9) with N-terminal Nluc-tag. B) and C) Measurement of total particle number (red line) by NTA (Nanoparticle Tracking Analysis), total protein in μg (green line) by BCA and total Nluc activity (blue line) in all single 500μL SEC fractions from F7 to F24 recovered from Nluc-CD63 or Nluc-CD9 supernatants. Shown data are from a single pilot experiment. D) and E) Western Blot analysis of SLC3A2/CD98, CD63, Syntenin, 14-3-3 and CD9 EV markers in EV-rich (F7-15) and protein-rich (F16-24) in Nluc-CD63 and Nluc-CD9 cells. Arrows indicate chimeric (Nluc-tagged) vs endogenous CD63 (D) or CD9 (E). F) and G) Representative TEM images showing CD63+ EVs (for Nluc-CD63) or CD9+ EVs (for Nluc-CD9) per μm2 isolated from 7*10^6 cells in F7-11 versus F12-24. Scale bar 0,5μm. Shown data are from a single pilot experiment. Arrowheads indicate EVs positive for CD63 staining (F) or CD9 staining (G).

Overall, we found that the majority of extracellular Nluc activity measured in Nluc-CD63 and Nluc-CD9 supernatants is associated to Nluc-tagged EVs, and only less than 30% could have a different origin (e.g. free proteins or small membrane fragments). Consequently, we concluded that measuring extracellular Nluc activity is a consistent and easy read-out of EV secretion compatible with high throughput screening procedures.

### Validation of the read-out for HCS screening by controlled manipulation of Nluc-CD63 and Nluc-CD9 secreting cells

Although validated in normal culture conditions, the use of our assay for screening a drug library brings additional constraints. One of the most obvious is the potential impact of some compounds contained in the drug library on cell viability, which would introduce a bias in the read-out of EV-associated Nluc release. Indeed, the MISEV2018 guidelines recommend to systematically report the level of cell viability in cultured cells producing EVs, since cell death may lead to release of both soluble and membrane bound structures such as apoptotic bodies [30]. Therefore, to investigate how cell death associated to potential compound toxicity could affect the measurement of Nluc activity in the supernatant, we treated Nluc-CD63 and Nluc-CD9 cells with Puromycin. Increasing doses of Puromycin led to a parallel increase of Nluc activity in the supernatant of both cell lines (Suppl. Fig.2A and 2B) and decrease of cell viability (quantified as number of Hoechst-labeled nuclei of the same cells, with dead cells characterized by round and bright nuclei). Thus, to avoid possible misinterpretations of our assay, we established a threshold of “acceptable” cell viability as around 85% of live cells, corresponding to 1.5μg/mL of Puromycin concentration, which increased extracellular Nluc activity by less than 30% (Suppl. Fig. 2A-B). To further challenge our assay, we measured Nluc activity in Nluc-CD63 and Nluc-CD9 cells treated with a drug known to increase exosome secretion, Bafilomycin A1[21][23][24]. As expected, Bafilomycin A1 treatment increased Nluc activity only in the supernatant of Nluc-CD63 cells (1.78 fold) but not in that of Nluc-CD9 cells (Suppl. Fig 2C-D). Importantly, in none of the cell lines this drug showed cell toxicity (Suppl. Fig. 2C-D).

**Figure 2.**
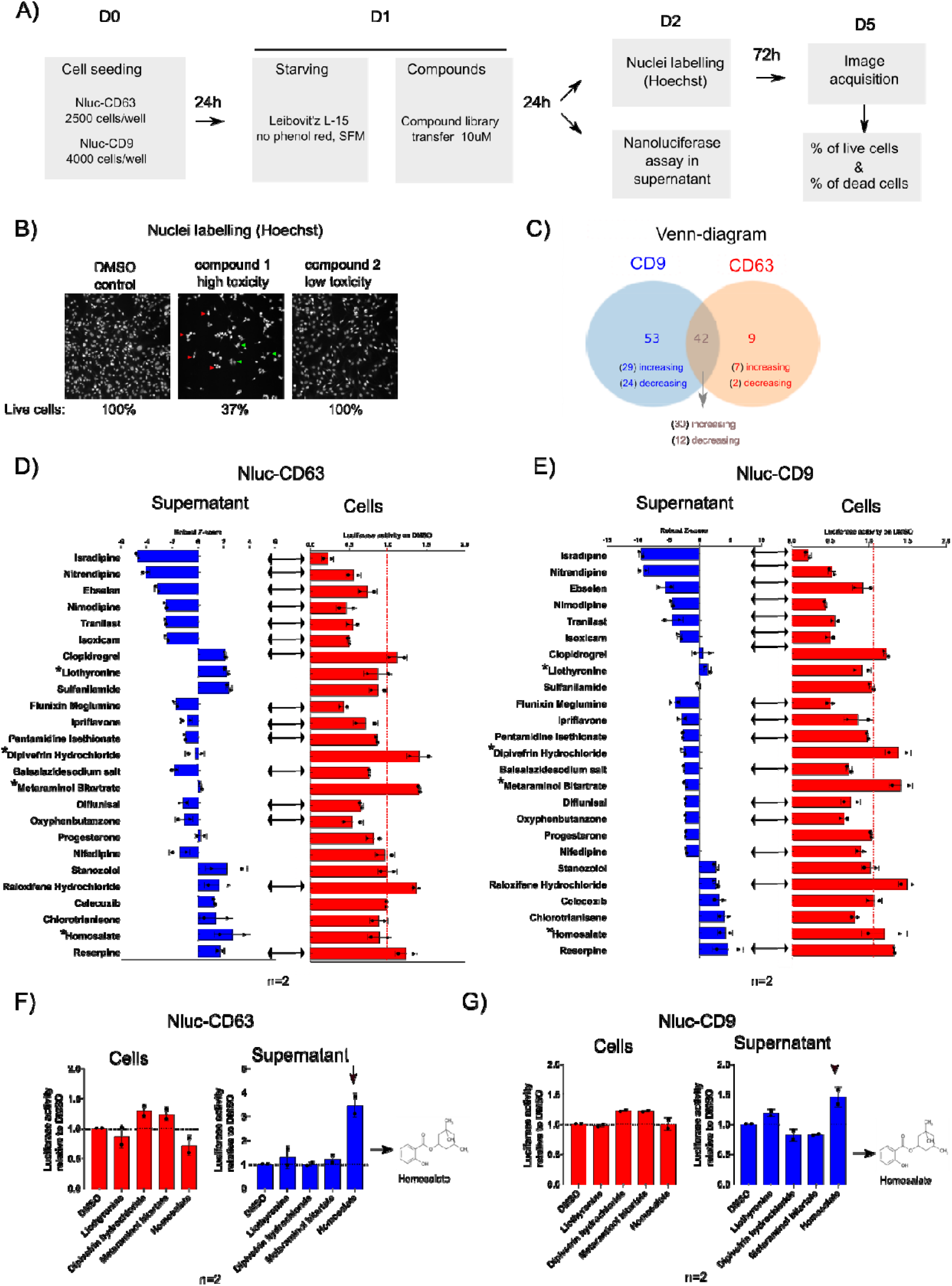
Identification of a drug increasing extracellular Nluc activity in Nluc-CD63 and Nluc-CD9 cells. A) Representative scheme of the screening protocol. SFM = serum-free medium. B) Representative example images of Hoechst nuclei staining for DMSO negative control (100% live cells), a high toxicity compound (=compound 1: 37% live cells) or a low toxicity compound (=compound 2: 100% live cells). Very round and bright nuclei are specific of dead cells. Total number of nuclei (i.e. total number of live + dead cells) and number of dead cell nuclei are counted, to calculate the actual number of live cells, in each condition as compared to the DMSO control. Green arrowheads: live cells; red arrowheads: dead cells. C) Venn-diagram summarizing the obtained screening results for two independent experiments. A total of 104 outlier compounds were identified. Among these, 53 only affected Nluc-CD9, 9 only affected Nluc-CD63 and 42 affected both cell lines. For each group, the number of increasing or decreasing outliers is reported. D) and E) Selection of 25 compounds from the 104 total outliers in Nluc-CD63 (D) and Nluc-CD9 (E) following criteria described in Suppl. Fig. 2F. Blue graphs: for each compound, extracellular Nluc activity intensity measured in the screening is reported as robust Z-score= [(compound value-median of (Ref pop))/(MADnc X 1.4826)], MAD=[median (|Ref pop-median (Ref pop)|)]. Increasing or decreasing hits were called according to the Threshold: |Robust Z score|>2 or <-2. Red graphs: for each compound, intracellular Nluc activity intensity was measured and reported as ratio on DMSO negative control. Data from two independent experiments are shown. Arrows between blue and red show non-selected compounds inducing the same trend of effect in cells and supernatants in both independent experiments, whereas * symbols indicate compounds selected for further validation. F) and G) Validation of four of the identified outliers. For Nluc-CD63 and Nluc-CD9 cells, intracellular (red) versus supernatant (blue) Nluc activity was measured after treatment with Lyothyronine, Dipivefrin Hydrochloride, Metaraminol Bitartrate and Homosalate. Data are expressed as ratio on DMSO negative control and are from two independent experiments.

This result shows that our assay is sensitive enough to reveal an increase of EV release regardless any possible cell death event. Moreover, it allows addressing the specificity of the effect towards distinct EV populations (CD63 and CD9 positive respectively).

### Identification of a drug increasing Nluc activity in the supernatant of Nluc-CD63 and Nluc-CD9 cells

With the established threshold to exclude effects of drug toxicity on Nluc activity, we next proceeded with the screening assay depicted in Fig. 2A to identify compounds able to affect the secretion of Nluc-CD63 EVs, Nluc-CD9 EVs, or both. We exposed Nluc-CD63 or Nluc-CD9 cells to a library of 1,280 FDA- and EMA-approved compounds (V3, Prestwick) at 10μM final concentration. We simultaneously measured Nluc activity in the cell supernatant, and cell viability by labelling the nucleus of the secreting cells with Hoechst, in order to monitor the toxicity of the administered compounds (Fig.2B). The screening was performed twice independently and for each drug, the level of extracellular Nluc activity was normalized by that of cells treated with DMSO as internal reference control. We expressed normalized extracellular Nluc activity as “Robust Z-score” (RZ-score, as described in materials and methods) and outlier compounds modulating EV secretion were defined by a RZ-score >2 or <-2. We thus identified 9 outlier compounds affecting Nluc-CD63, 53 outlier compounds affecting Nluc-CD9 and 42 outlier compounds affecting both (Fig. 2C, Suppl. Fig. 2E). To select the hits to use for further validation, we thus applied the cell death criteria described in Suppl. Fig. 2F. We discarded 79 drugs resulting in less than 80-85% of live Nluc-CD63 or Nluc-CD9-cells. We selected 25 among the 104 total outlier compounds identified in the screening (see graphs in blue in Fig. 2D and 2E) for which we secondarily measured the effect on intracellular Nluc activity (see graphs in red in Fig. 2D and 2E). As we were seeking for drugs affecting specifically the release of EV-associated Nluc and not expression of the Nluc-fused molecules nor Nluc enzymatic activity, we discarded all the compounds predicted to increase or decrease Nluc activity in the supernatant which were showing the same effect on intracellular Nluc activity (arrows in Fig. 2D-E). Thus, we selected a group of four compounds modulating EV secretion: two that decreased (Dipivefrin hydrochloride, Metaraminol bitartrate) versus two that increased (Liothyronine, Homosalate) EV secretion in at least one of the two cell lines. We then re-validated them in both cell lines for intracellular or supernatant Nluc activity (Fig. 2F and 2G). Among all the selected hits, Homosalate increased in the strongest and most reproducible manner the release of Nluc activity by both cell lines, thus we focused on this drug for further validation (Fig. 2F and 2G).

### Homosalate increases EV secretion enriching a population of SLC3A2/CD98-positive EVs

To confirm the effect of Homosalate using classical EV isolation and characterization methods, we treated MDA-MB-231 parental cells with Homosalate and isolated EVs using SEC. We confirmed no detrimental effect of Homosalate on cell viability by Trypan Blue staining (Fig.3A). Strikingly, we could measure a significant increase in particle number by NTA both in the total cell supernatant (input) and the EV-rich F7-11 fractions (Fig.3B). We confirmed by TEM that Homosalate increased the number of CD63- and of CD9-positive EVs (Fig. 3C). We next analyzed the expression of EV markers in isolated EVs (F7-11) by Western Blot. EV samples isolated from the same number of secreting cells were loaded on the gels (Fig.3D). We observed an increase of EV makers SLC3A2/CD98, CD63, Syntenin, CD9 and CD81 (Fig 3D) whereas no major changes were observed in the total lysates (CL) (Fig 3D). Taken together, these data showed that Homosalate is an increaser of EV secretion in parental MDA-MB-231. Importantly, Homosalate also increased EV (or at least particle) release in two other tumor cell lines, HeLa and MCF7 with minimal impact on cell viability, and in another one, Jurkat, but with a significant toxicity that, for this cell line, could participate in the observed increased EV release (Suppl. Fig. 3A) (Suppl. Fig. 3B). To determine whether, besides increasing the total amount of released EVs, Homosalate also affected their composition, we next analyzed EV secreted by Homosalate-treated MDA-MB-231 parental cells by WB loading the same number of particles (Fig.3E). We thus observed that only SLC3A2/CD98 and to a minor extent CD9 were increased (Fig. 3E). These data suggested that Homosalate primarily increased a population of EVs enriched in SLC3A2/CD98 and CD9. Consistently, our group previously showed that SLC3A2/CD98 is present on distinct CD63- or CD9-positive EVs populations secreted by HeLa cells, and is particularly enriched in the latter under certain conditions [18]. To characterize EV subpopulations secreted by MDA-MB-231, we performed an EV co-immunoprecipitation assay using anti-CD63- or anti-CD9-coated beads in SEC-isolated EVs. Consistent with [18], SLC3A2/CD98 co-immunoprecipitated with both CD63 and CD9, but the percent of SLC3A2/CD98 that co-immunoprecipitated with CD9 (97%) was higher than the percent co-immunoprecipitated with CD63 (86%) (Fig. 3F). Similarly, we observed that only 83% of CD9 co-immunoprecipitated with CD63 (Fig.3F). Finally, only 63% of CD63 co-immunoprecipitated with CD9 (Fig. 3F). These findings suggest that MDA-MB-231 secrete heterogeneous subpopulations of EVs characterized by different combinations of CD63, CD9 and SLC3A2/CD98, with a predominant sub-population of EVs characterized by simultaneous expression of SLC3A2/CD98 and CD9 with CD63 and two minor sub-populations characterized respectively by co-expression of SLC3A2/CD98 and CD9 without CD63, or by CD63 alone. Taken together, these data indicate that Homosalate not only induces a sustained release of bulk EVs (Fig. 3B-C-D), but also increases a subpopulation of EVs double positive for SLC3A2/CD98 and CD9 (Fig. 3F) which could be more sensitive to its effect (Fig. 3E).

**Figure 3.**
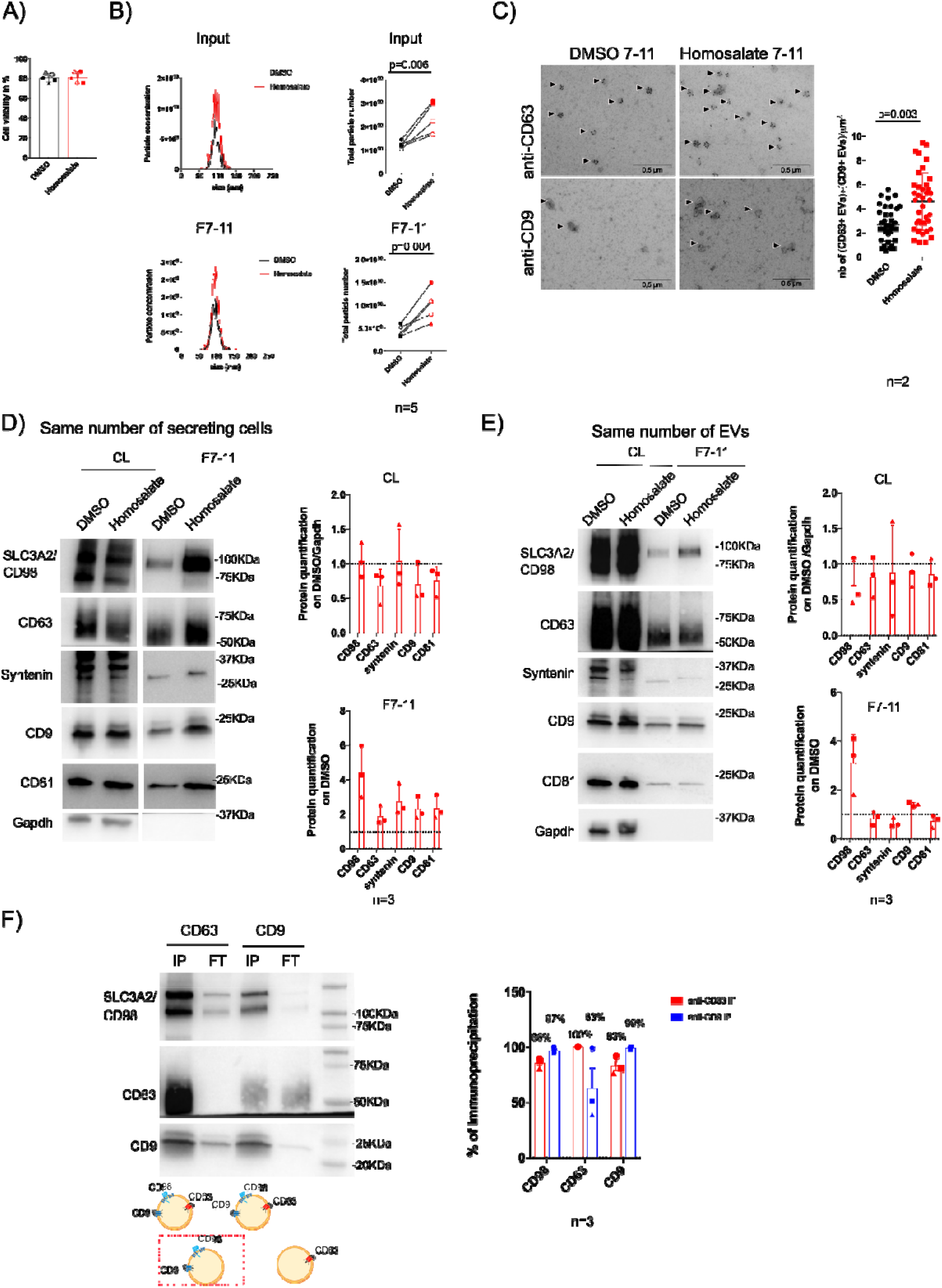
Homosalate increases EV secretion enriching a population of SLC3A2/CD98-positive EVs. A) Quantification of cell viability after Homosalate treatment in MDA-MB-231 parental cells. DMSO or Homosalate treated cells were counted after collection of conditioned media, using Trypan Blue as a reporter of cell death. Cell viability is expressed in percentage. Data from five independent experiments are shown. B) Quantification of EVs induced by treatment with Homosalate. Left panel: one representative graph showing particle concentration/cm^3^ versus particle size measured by NTA in inputs (total conditioned medium) and F7-11 (EV-rich SEC fractions) from 27*10^6 DMSO or Homosalate treated cells. Right panel: Graphs show total particle number secreted from 27*10^6 cells measured by NTA in DMSO or Homosalate treated cells for inputs and SEC F7-11, from five independent experiments. Paired parametric t-test p=0.006; p=0.004. C) Representative TEM images showing (CD63+) or (CD9+) EVs in F7-11 released by 2.7*10^6 DMSO or Homosalate treated cells. Arrowheads indicate EVs positive for CD63 staining (above) or CD9 staining (below). Graphs show quantification of the number (=nb) of (CD63+EVs) + (CD9+ EVs) per μm^2^. Scale bar 0.5μm. Data from two independent experiments are shown, each dot represents EVs counted in one field (DMSO: 20 dots for replicate 1, 18 dots for replicate 2; Homosalate: 20 dots for replicate 1, 18 dots for replicate 2). Mann-Whitney test p=0.003. D) Western Blot analysis of EV markers SLC3A2/CD98, CD63, Syntenin, CD81 and CD9 released by cells treated with DMSO or Homosalate after gel loading with same number of secreting cells. Gapdh was used as normalizer for cell lysates (CL). CL from the equivalent of 200 000 cells were loaded, F7-11 from the equivalent of 2.7*10^6 secreting cells were loaded. Graphs show protein signal quantifications normalized first on Gapdh and then on DMSO for CL or normalized on DMSO for F7-11. Data from three independent experiments are shown. E) Western Blot analysis of EV markers SLC3A2/CD98, CD63, Syntenin, CD81 and CD9 released by cells treated with DMSO or Homosalate, after gel loading with same numbers of particles. Gapdh was used as normalizer for CL. CL from the equivalent of 200 000 cells or an amount corresponding to 4*10^8 particles for F7-11 were loaded. Graphs show protein signal quantifications normalized first on Gapdh and then on DMSO for CL or normalized on DMSO for F7-11. Data from three independent experiments are shown. F) Western Blot analysis of EVs immunoprecipitated from F7-11 and schemes of the EVs recovered. The equivalent of 1*10^9 particles in F7-11 were immunoprecipitated with anti-CD63- or anti-CD9-coated beads (IP) and loaded side-by-side with the corresponding unbound EVs contained in the flow-through (FT). Blots were incubated with SLC3A2/CD98, CD63 or CD9 antibodies. Percent of immunoprecipitation displayed in the right panel graph was calculated as signal for SLC3A2/CD98, CD63, CD9 present in CD63 or CD9 IP divided by the total signal for the protein in the same IP: IP/(IP+FT). Data from three independent experiments are shown.

### Homosalate and Bafilomycin A1 induce the release of EV subpopulations generated in distinct subcellular compartments

Bafilomycin A1 is a known specific increaser of release of exosomes [23][24][25]. We asked whether Homosalate showed the same specificity by comparing the effects of the two drugs. As expected, neither Homosalate nor Bafilomycin A1 increased cell death in parental MDA-MB-231 (Fig. 4A) and both increased the number of particles by NTA (Fig. 4B). Then, when we analyzed the expression of EV markers by Western Blot normalized by same number of particles, we observed that Homosalate reproducibly increased the expression of SLC3A2/CD98 (in 4/5 biological replicates) (Fig. 4C). Conversely, CD63 release in EVs was increased by Bafilomycin A1 in 4/5 replicates, suggesting a specific effect of this drug on CD63, consistent with previous observations [18][21][24] (Suppl. Fig. 2C). Lamp-1 is a marker of late endosomal compartments [31]. As expected, we found an increase of Lamp-1 after treatment with Bafilomycin A1 in all 5 replicates (Fig. 4C). Thus, the specific increase of Lamp-1 after Bafilomycin A1, but not Homosalate treatment suggests that the two drugs are acting on distinct subcellular compartments. To better define the subcellular origin of EVs induced by Homosalate or Bafilomycin A1, we analyzed the intracellular localization of CD63, CD9 and SLC3A2/CD98 markers by immunofluorescence. As shown previously for HeLa cells [18], we observed that CD63 was localized mostly in intracellular compartments in DMSO-treated negative control, whereas CD9 was mainly localized at the plasma membrane and in few intracellular compartments (Fig. 4D, left panel). Interestingly, SLC3A2/CD98 co-localized with CD9. After treatment with Bafilomycin A1 we found an increased number of large CD63-positive perinuclear compartments and CD9 and SLC3A2/CD98 appeared to be more accumulated intracellularly (Fig 4D, right panel). Different to this, we observed that after Homosalate treatment none of the analyzed markers were accumulated intracellularly and instead, SLC3A2/CD98 and CD9, but not CD63, were enriched at the plasma membrane (Fig. 4D, middle panel). Co-localization measurements of the analyzed markers revealed that the amount of SLC3A2/CD98 co-localizing with CD9 in the DMSO control was around 80%, higher than the one co-localizing with CD63 (around 50%) (Fig. 4E). This observation corroborates the data shown in Fig. 3F, in which SLC3A2/CD98 preferentially co-immunoprecipitated with CD9 than with CD63. Co-localization of CD63 and CD9 was also significant (70%). Homosalate did not change the amount of SLC3A2/CD98 nor of CD63 co-localizing with CD9, whereas Bafilomycin A1 increased both to respectively 85% and 80% (Fig. 4D). By contrast, Homosalate treatment reduced the amount of SLC3A2/CD98 co-localizing with CD63 to 30%, whereas Bafilomycin A1 treatment increased it to 70%, indicating that this drug causes the two markers to be more accumulated in late endosomal compartments (Fig. 4D).

**Figure 4.**
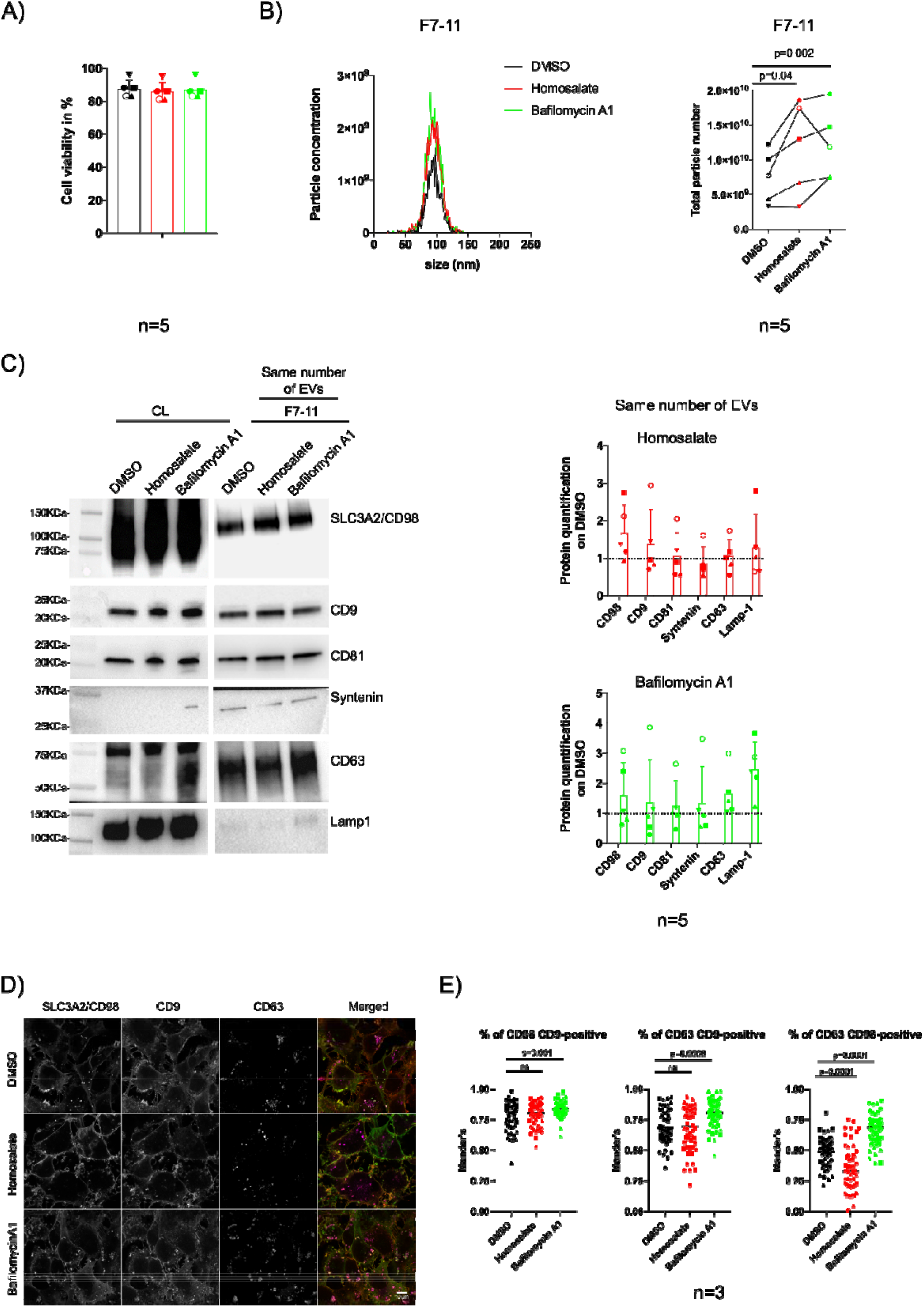
Homosalate increases the secretion of plasma membrane derived EVs. A) Quantification of cell viability after Homosalate and Bafilomycin A1 treatment in MDA-MB-231 parental cells. DMSO, Homosalate or Bafilomycin A1 treated cells were counted after conditioned media collection, using Trypan Blue as a reporter of cell death. Cell viability is expressed in percentage. Data from five independent experiments are shown. B) Quantification of EVs induced by treatment with Homosalate and Bafilomycin A1 in MDA-MB-231 parental cells. Left panel: Graphs show particle concentration/cm^3^ versus particle size measured by NTA in F7-11 (EV-rich SEC fractions) from 50*10^6 DMSO, Homosalate or Bafilomycin A1 treated cells. Right panel: Graphs show total particle number secreted from 50*10^6 cells measured by NTA in DMSO, Homosalate or Bafilomycin A1 treated cells for SEC F7-11. Data from five independent experiments are shown. Paired parametric t-test p=0.04; p=0.002. C) Western Blot analysis of EV markers SLC3A2/CD98, CD9, CD81, Syntenin, CD63 and Lamp-1 released after treatment with DMSO, Homosalate or Bafilomycin A1 after gel loading with same number of particles. CL from the equivalent of 200 000 cells or an amount corresponding to 4*10^8 particles from F7-11 were loaded. Graphs show protein signal quantifications normalized to DMSO for F7-11 after treatment with Homosalate (above) or Bafilomycin A1 (below). Data from five independent experiments are shown. D) Immunofluorescence of SLC3A2/CD98, CD63 and CD9 in MDA-MB-231 treated with DMSO, Homosalate or Bafilomycin A1. E) Graphs show Mander’s correlation coefficients for CD98-CD9, CD63-CD9 or CD63-CD98 co-localization expressed as percentage. Ordinary one way Anova, multiple comparison test CD98-CD9: ns or p=0.001; CD63-CD9: ns or p=0.0008; CD63-CD98 p=0.0001. Data from three independent experiments are shown, each dot represents one counted cell (DMSO: 25 dots for replicate 1, 10 dots for replicate 2, 19 dots for replicate 3; Homosalate: 25 dots for replicate 1, 12 dots for replicate 2, 21 dots for replicate 3; Bafilomycin A1: 21 dots for replicate 1, 16 dots for replicate 2, 22 dots for replicate 3).

Taken together, these data suggest that Homosalate and Bafilomycin A1 induce the release of distinct EV subpopulations. Bafilomycin A1 acts on EVs originating into intracellular, late endosomal compartments (=exosomes) enriched with CD63 and with an intracellular pool of SLC3A2/CD98 and CD9, whereas Homosalate mostly acts on a subpopulation of EVs enriched in SLC3A2/CD98 and CD9, which could rather originate directly at the PM (Figure 4D, middle panel) (=ectosomes) as previously described in HeLa [18].

### Homosalate, but non Bafilomycin derived EVs induce resistance to anoikis

Since Homosalate and Bafilomycin A1 induce the release of EVs derived from different subcellular compartments (Fig. 4D), we wondered whether Homosalate- or Bafilomycin A1-induced EVs could display different functional properties. Homosalate has been shown to increase in vitro migratory and invasive properties of several breast cancer cell lines when administered at low concentrations (≥100nM) during several weeks [32] and to be highly toxic at certain concentrations for luminal breast cancer MCF7 cells [33]. Based on this knowledge, we wondered whether the EVs generated by MDA-MB-231 cells upon Homosalate treatment induced functional consequences when applied to recipient tumor cells. First, after isolation of EVs (F7-11) from Homosalate- or Bafilomycin A1-treated MDA-MB-231, we administered 5×10^8^ of each type of EVs to MCF7 during 1h and then we followed their behaviour in real-time during 50h using the XCellIgence system to measure continuously adhesion and number of cells, as a proxy for cell growth. As shown in Fig. 5A, the growth rate of MCF7 remained constant, in absence (=No EVs) or presence of either types of EVs.

**Figure 5.**
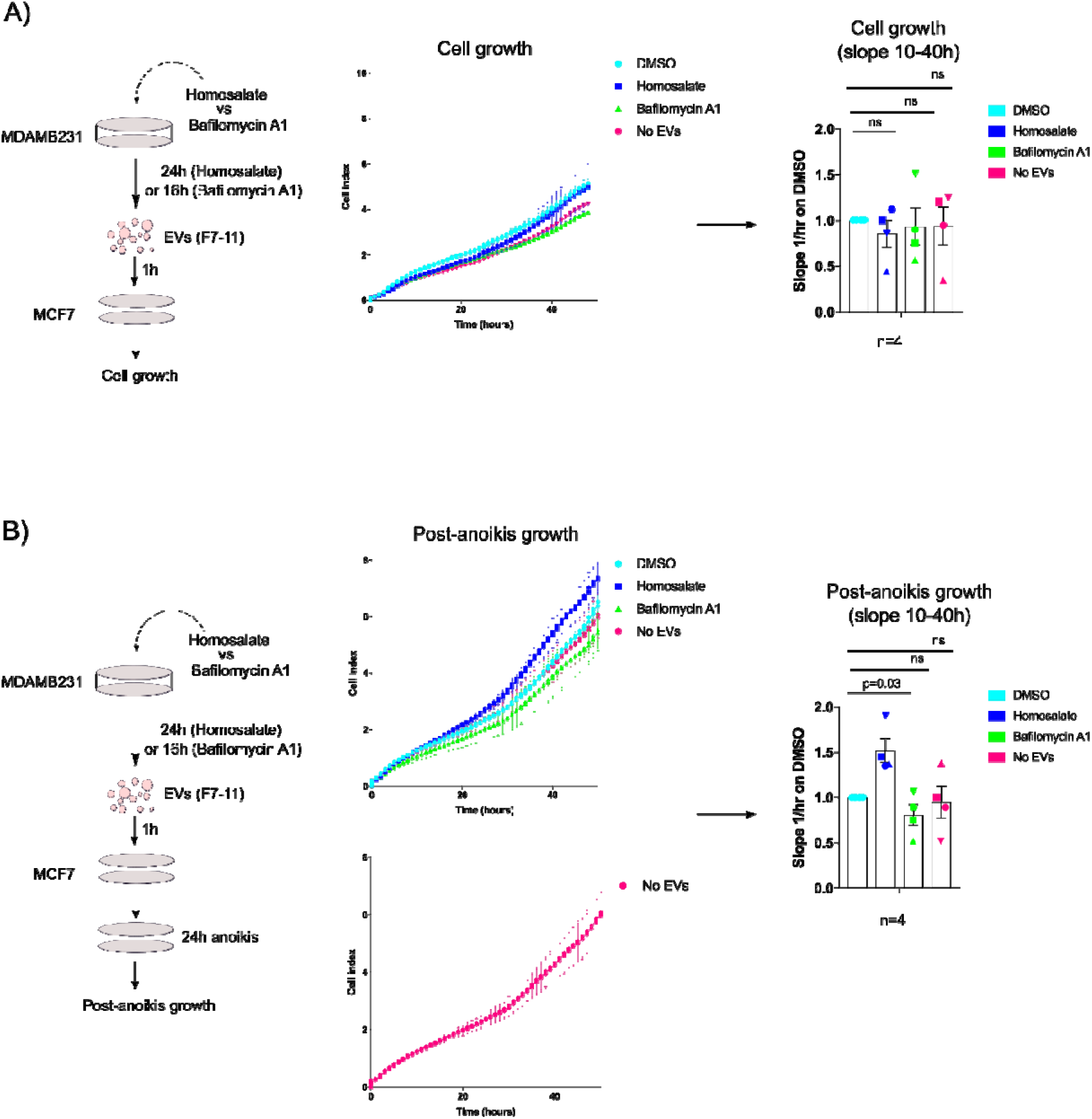
Homosalate, but non Bafilomycin A1 derived EVs induce resistance to anoikis. A) Left panel: Representative scheme of EV uptake experiment in MCF7 cells. Middle panel: Representative graph of MCF7 real time adhesion and proliferation 1 hour after uptake of EVs from DMSO, Homosalate or Bafilomycin A1 treated MDA-MB-231. Measurements were programmed every 30 minutes for a total time of 50 hours in XCELLigence device. Right panel: graph showing quantification of slopes in the range 10-40 hours. Data are expressed as ratio on DMSO negative control and are obtained from four independent experiments. Ordinary one-way Anova test non-significant (ns). B) Left panel: Representative scheme of EV uptake followed by anoikis assay in MCF7 cells. Middle panel: Representative graph of MCF7 real time adhesion and proliferation 1 hour after uptake of EVs from DMSO, Homosalate or Bafilomycin A1 treated MDA-MB-231 and 24 hours of anoikis assay. Measurements were programmed every 30 minutes for a total time of 50 hours in XCELLigence device. Right panel: graph showing quantification of slopes in the range 10-40 hours. Data are expressed as ratio on DMSO negative control and are obtained from four independent experiments. Ordinary one-way Anova test p=0.03 for Homosalate and non-significant (=ns) for the others.

We then reasoned that the migratory and invasive behaviour modulated by Homosalate could more rely on cell-cell or cell-ECM adhesion properties. Deprivation of extracellular matrix support can induce a particular type of apoptosis called “anoikis” [34]. Both tumor and non-tumor cells of epithelial origin can resist to anoikis by enforcing cell-cell contacts and forming large cell aggregates [35] [36] or can become more resistant to anoikis under certain conditions [37]. In several cases, resistance to anoikis has been considered as hallmark of increased tumor aggressiveness [38][39][40]. To evaluate whether EVs could affect the capability of MCF7 to grow in absence of cell-matrix contacts, we fed MCF7 cells with a fixed number of EVs for 1h, before subjecting them to an anoikis assay, in which cells are grown for 24h in the absence of cell-matrix contacts [37]. Post-anoikis cell growth was followed in real time over 50 hours. MCF7 cells fed with EV derived from Homosalate-treated cells survived and proliferated more compared to EV from Bafilomycin A1-treated or control cells (Fig. 5B). We proved that this effect was not merely due to EV treatment, because the “No EVs” control gave results similar to DMSO or Bafilomycin A1 EVs (Fig. 5B).

Taken together, these data indicated that EVs generated by MDA-MB-231 cells upon treatment with Homosalate are likely derived from PM and conferred to MCF7 cells enhanced capability to grow in harsh conditions i.e. in the absence of a supportive extracellular matrix, which could increase their tumorigenic potential. Conversely, MVB-derived EVs induced by Bafilomycin A1 were not able to confer to MCF7 the same properties.

In conclusion, we show that specific drug treatments differentially affect not only the release and the composition of EVs but most importantly also their function.

## Discussion

In the work presented here, we 1) established tools and a robotized process to quantify EVs bearing either one of two different markers, in a few microliters of cell conditioned medium, 2) successfully screened a bank of over 1200 health agency-approved drugs for effect on EV release, 3) identified Homosalate as a potent drug increasing release of EVs likely originated from PM, 4) showed that EVs released upon Homosalate treatment promote cell survival in non-attachment conditions, not observed with Bafilomycin A1-induced EVs.

In the literature, the number of chemical compounds affecting specifically release of given EV subtypes is still limited. For example, the drugs GW4869 [41] or Manumycin A [42] have been proposed to decrease exosome release by ESCRT-independent or -dependent mechanisms, respectively. However, no direct proof was provided that these drugs did not affect the other EV types. A phospholipase D2 inhibitor, CAY10594, inhibits specifically secretion of Syntenin-containing exosomes formed by an ARF6-dependent pathway [43], but ARF6 is also involved in ectosome release by other cells [44]. Recently, screening of a Syntenin-PDZ domain-focused fragment library led to the identification of a small molecule decreasing specifically the release of Syntenin-containing exosomes [45]. The compound Y27632 decreases PM-derived ectosome release by targeting ROCK1 and ROCK2 kinases [46]. Conversely, very few compounds have been shown to increase EV secretion. Bafilomycin A1 and other inhibitors of V-ATPase activity [21] affect primarily exosome secretion. Ionomycin, which increases intracellular Ca^2+^ levels [47], increases release of exosomes but also of other types of EVs, and can induce cell death. Therefore, screenings of small molecule libraries designed at distinguishing effects on EV subtypes and eliminating death-inducing confounding factors represent novel valuable approaches to identify specific modulators of EV release and composition.

We choose Nluc to establish a miniaturized and robotized process of EV quantification because, a priori, Nluc is an excellent tool to be used as reporter of biological activity. Indeed, this enzyme is considerably smaller (its size is ∼19kDa) than the traditionally used luciferase enzymes (e.g. Renilla or Firefly luciferases) and has the minimum impact on the topology of tagged proteins. Most importantly, Nluc enzymatic activity is up to 100 folds more efficient than that of other luciferase enzymes [28]. Nevertheless, Nluc is sensitive to the action of some chemical compounds which can inhibit its enzymatic activity [48][49]. Here, as previously reported [50], we successfully used Nluc to tag either CD63- or CD9-positive EVs and follow their secretion directly in the supernatant of cells (Fig.1). When we performed a HCS of chemical compounds, we found several hits affecting Nluc activity in the supernatant of cells (Fig.2D-E, Suppl. Fig. 2E). The specificity of action of these hits on EV release was carefully assessed using intracellular Nluc activity as control, to exclude effects on the expression of the fusion reporters (CD63, CD9) or on Nluc activity itself. Among the drugs decreasing extracellular Nluc, Isradipine had been described to inhibit Nluc enzymatic activity [49] (Fig.2D-E). Other compounds of the same family containing a phenyl-1,4-dihydropyridine core, like Nitrendipine, Nimodipine or Nifedipine, all had a similar effect (Fig. 2D-E), suggesting that they directly inhibited Nluc activity, rather than inducing a decrease in EV release (Fig. 2D-E). To increase the stringency of our screening procedure, we thus decided to exclude from further validation compounds that affected intra- and extracellular Nluc activity in the same manner (i.e. both decreasing or both increasing, see arrows in Fig. 2D-E). We also identified cell death as a critical parameter to be considered as it could increase leakage from the cell or the release of other membrane-bound structures (e.g. apoptotic bodies) [30] (Suppl. Fig. 2A-B). Therefore, an important aspect of our screen was the systematic measurement of cell viability of drug-exposed cells. Finally, it should be mentioned that, because ∼ 30% of Nluc activity measured in the supernatant comes from elements other than EVs (e.g. free proteins, debris, etc.) (Fig. 1B-C), drugs affecting release of EVs by less than 30% could not be reliably identified.

A low-throughput drug screening using Nluc activity as a read-out was previously successfully used [21]. The authors screened selected drugs and identified Bafilomycin A1 (and other compounds acting on V-ATPases) as increasers of Nluc-CD63 EV secretion [21]. We could confirm that Bafilomycin A1 increased extracellular Nluc activity in Nluc-CD63 but not Nluc-CD9 cells (Suppl. Fig. 2C-D), highlighting a differential effect of this MVB-modulating drug on CD63 or CD9 positive EV secretion. In another study, the use of a dual reporter system of CD63-Turbo-Luciferase-CD9-Emerald-Green allowed to identify several compounds regulating the secretion of CD63-CD9 double positive EVs by immune cells [51]. Turbo-Luciferase could be particularly suitable for high-throughput screenings because it is less sensitive to chemical compound interference [52].

Our screening allowed the identification of Homosalate as robust increaser of EV secretion in MDA-MB-231 parental cells and in other tumor cell lines (Fig.3, Suppl. Fig.3). Homosalate specifically increases the secretion of EVs characterized by the simultaneous expression of SLC3A2/CD98 and, to a minor extent, of CD9 (Fig.3E). In support of this, we observed that in MDA-MB-231 cells up to 97% of SLC3A2/CD98 co-immunoprecipitated with CD9, but only 86% co-immunoprecipitated with CD63 (Fig.3F). These observations are consistent with previous findings from our group, showing that SLC3A2/CD98 was co-expressed with CD9 more than with CD63 on EVs secreted by HeLa cells, and that CD9-positive EVs were mainly generated at the PM (=ectosomes), whereas CD63-positive EVs were mainly generated in MVBs (=exosomes) [18]. Consistently, in our MDA-MB-231 cellular model in control conditions (=DMSO), CD63 was mainly localized in intracellular perinuclear compartments, whereas CD9 and SLC3A2/CD98 were more (but not exclusively) localized at the PM (Fig.4D). Indeed, in these conditions, we quantified 80% of SLC3A2/CD98 co-localizing with CD9, but only 50% of CD63 co-localizing with SL3CA2/CD98 (Fig.4E). Consistent with our previous observations [18], in MDA-MB-231, the analyzed EV markers were not restricted to a single compartment (i.e. only PM or only MVB), as for SLC3A2/CD98 or CD9 which are detected in few intracellular compartments besides PM (Fig.4D). We compared the effect of Homosalate to that of Bafilomycin A1, which acts on MVBs and increases exosome release. Consistent with this, Bafilomycin A1 modulated CD63 more than SLC3A2/CD98 in Western Blot, while Homosalate had an opposite effect (Fig.4C). Microscopy analysis showed that Homosalate increased membrane staining of SLC3A2/CD98 and CD9 but overall did not change the percentage of SLC3A2/CD98 or CD63 co-localizing with CD9. On the contrary, it decreased to 30% the amount of CD63 co-localizing with SLC3A2/CD98 (Fig.4E). Bafilomycin A1 seemed to act in a different manner, because it increased the accumulation of CD63, CD9 and SLC3A2/CD98 in intracellular compartments (Fig.4D), and greatly increased the co-localization of CD9 or SLC3A2/CD98 with CD63 (Fig.4E). This suggests that Bafilomycin A1 acts on the intracellular pools of these proteins which will be subsequently co-expressed on secreted exosomes, as opposed to Homosalate which would act on the membrane pools of SLC3A2/CD98 and CD9 to give rise to ectosomes.

Homosalate was chosen because of its prominent effect on extracellular Nluc activity both in screening and in further validation steps. Other compounds also had relevant effects on EVs expressing CD63, CD9 or both. For example, even though their effect on extracellular Nluc activity was not as striking as that of Homosalate, Metaraminol Bitartrate or Dipivefrin Hydrochloride could be promising specific inhibitors of CD9-containing EVs release (Fig.2F-G), since they induced a consistent decrease of extracellular Nluc activity in screening and further validation steps (Fig.2D-E-F-G). Regular EV quantification and characterization analyses are still needed to validate these drugs as novel EV inhibitors.

Homosalate is an anti-inflammatory drug commonly used as a chemical UV-screen in sun lotions [53][54]. However, it displays estrogen-like properties and is suspected to be a potential endocrine disruptor dangerous for health [55]. Here we identified an unsuspected action of Homosalate, i.e. increasing EV release from different tumor cell lines. We demonstrated that in MDA-MB-231 triple negative breast cancer cells, EVs induced by Homosalate are generated in a distinct subcellular compartment compared to those induced by Bafilomycin A1. Not last, we demonstrated that EVs secreted by MDA-MB-231 upon treatment with Homosalate are endowed with a particular function: making other recipient tumor cell lines (i.e. MCF7 luminal breast cancer cells) more resistant to anoikis. Resistance to anoikis (i.e. death occurring due to loss of cell matrix contacts) can be a hallmark of increased tumor aggressiveness [38][39][40], although it can be also a property of some non-tumor epithelial cells [35]. The safety of Homosalate in cosmetics is still a matter of debate [56]. Although our findings must be completed by additional studies on effective capacity of Homosalate to contribute to cancer progression, they could worsen the concern about using this compound in certain skin-care products.

## Materials and methods

We have submitted all relevant data of our experiments to the EV-TRACK knowledgebase [57], with the following accession number (EV-TRACK-ID) for reviewers: EV210281.

### Cell culture and transfection

MDA-MB-231, MCF7, and HeLa were cultured in our laboratory for the last 20 years, after initial obtention from the American Type Culture collection (ATCC). They were validated by short tandem repeat (STR) sequencing in 2018. Jurkat E6-1 cells were obtained from the America Type Culture Collection (ATCC). MDA-MB-231, MCF7 and HeLa were cultured in Dulbecco’s modified Eagle’s medium (DMEM-Glutamax™, Gibco), with 10% of Fetal Calf Serum (FCS, Gibco), 100U/mL penicillin and 100μg/mL streptomycin (Gibco). Jurkat were cultured in Roswell Park Memorial Institute 1640 medium (RPMI-1640-Glutamax™, Gibco), with 10% of Fetal Calf Serum (FCS, Gibco), 100U/mL penicillin and 100μg/mL streptomycin (Gibco). Cell lines were grown at 37 °C, under 5% CO2, in humidified incubators and routinely tested using Myco detection Kit (Eurofins) for mycoplasma contamination. Only mycoplasma negative cells were used for experiments. MDA-MB-231 bulk or stable populations overexpressing Nluc-CD63 or Nluc-CD9 constructs were obtained using an electroporation-based transfection protocol optimized for this cell line (Amaxa® Cell Line Nucleofector® Kit V, Lonza). Briefly, 1*10^6 cells were harvested by trypsinization and were mixed with 100μL of room temperature reconstituted Nucleofector® solution combined with 2μg of the DNA construct of interest. The solution was transferred into a certified cuvette and cell electroporation was conducted using Nucleofector® program X-013 (Amaxa® Nucleofector®, Lonza). Electroporated cells were immediately resuspended in pre-warmed phenol-free Leibovitz’s-L-15 medium (Fisher) with 10% of FCS and seeded in a 24 well plate. After 24 hours, cells were harvested by trypsinization and resuspended in DMEM-Glutamax medium with 10% of FCS, 100U/mL penicillin, 100μg/mL streptomycin and 2mg/mL of geneticin as selection antibiotic. A portion of the resuspension volume corresponding to 1/3 was seeded in 10 cm dishes for low density culture to obtain clonal populations and a portion corresponding to 1/10 was seeded in a 24 well plate to obtain a bulk population. Clonal populations resistant to geneticin selection appeared after ∼20 days of culture and were collected manually using cloning discs (Sciencewere® cloning discs, Sigma-Aldrich). Collected clones were transferred in 96 well plates and once they reached confluence they were further selected by mean of Nluc activity measurement (Nano-Glo® Luciferase Assay System, Promega, see details in sections below) in supernatants or in cells.

### Plasmids

Nluc sequence was PCR derived using the indicated primers (Forward: `ATTACTACCGGTATGGTCTTCACACTCGAAGATTTC; Reverse:` ATTACTCTCGAGCGCCAGAATGCGTTCGCACAG) from a Nluc construct (kind gift of Michael Boutros). Nluc-CD63 construct was obtained by removing RFP sequence from RFP-CD63 (kind gift of Walther Mothes) using AgeI and XhoI restriction enzymes (New England Biolabs) and by replacing it with Nluc sequence using the same enzymes. Nluc-CD9 construct was obtained as follows: CD9 sequence was first PCR derived from tdTomato-CD9-10 construct (kind gift of Michael Davidson) using the indicated primers (Forward: CTCAAGCTTCCCCGGTCAAAGGAGGCA; Reverse: ATCCGCAGGAACCGCGAGATGGTCTAG). Then, a Renilla-luciferase-HSP70 construct previously obtained in the lab by removing GFP sequence from GFP-HSP70 construct (Addgene, #1525) with XhoI and SpeI restriction enzymes (New England Biolabs) was used as intermediate. Briefly, HSP70 sequence was removed from Renilla-luciferase-HSP70 and was replaced by CD9 sequence using XhoI and SpeI restriction enzymes. Finally, Renilla-luciferase sequence was removed using AgeI and XhoI restriction enzymes and was replaced by Nluc sequence using the same enzymes.

### Nanoluciferase detection assay

Nano-Glo® Luciferase Assay System (Promega) was used to quantify Nluc activity both in cells and in supernatants of Nluc-CD63 and Nluc-CD9 cells. Experiments were conducted in 96 well plates (Fig. 2D and E), 384 well plates (Fig. 2D and E, Suppl. Fig. 2A, B, C and D) or 24 well plates (Fig. 2F and G). First, cells were seeded in the various multi-well formats according to their different growth rates: 96 well plates: [16,000 Nluc-CD63 and 26,000 Nluc-CD9]; 384-well plates: [2,500 Nluc-CD63; 4000 Nluc-CD9]; 24-well plates: [101000 Nluc-CD63; 165000 Nluc-CD9]. After 24 hours, cells were washed 1 X with phenol free, serum free Leibovitz’s L-15 medium and the same medium was added on cells (for 96-well plates: 100μL, for 384-well plates: 50μL, for 24-well plates: 500μl). When required by the experiment (as for Fig. 2 and Suppl. Fig.2), drugs were added at this step to the phenol red free, serum free Leibovit’z L-15 incubated on cells (DMSO 0,1%, Bafilomycin A1 100nM, Puromycin 1;1.5;2;3μg/mL). To measure Nluc activity in the supernatant of cells, we proceeded as follows: to avoid to disturb adherent cells, only a fraction of cell conditioned medium was retrieved (for 96-well plates: 80μL, for 384-well plates: 40μL and for 24-well plates: 50μL). Then, collected supernatants were centrifuged at 350g for 10 minutes at RT to eliminate dead cells and debris. To prevent disruption of cell pellet, only a fraction of the resulting centrifuged supernatant (for 96 well plates: 60 μL, for 384 well plates: 30 μL, for 24 well plates: 40 μL) was collected and transferred to 96 well white plates (Corning® #3912). For Nluc activity measurement, the reagent of Nano-Glo® Luciferase Assay System was reconstituted according to manufacturer’s instructions and was added to the supernatant at a 1:6 ratio. When performed, the measurement of intracellular Nluc activity was done as follows: cells were first washed 1X with phenol free, serum free Leibovitz’s L-15 medium leaving a residual of 25 μL of medium, and then Nano-Glo® Luciferase Assay reagent was added at a 1:4 ratio. Luminescence activity was read using iD3 SpectraMax microplate reader (Molecular Devices, California, USA) or Centro LB 960 microplate luminometer (Berthold, Germany). For Nluc activity measurement during drug screening, a specific protocol is fully described in the section below.

### Screening of the drug library

#### Cell seeding and library addition

Cells were amplified over a week before the screening step. For cell passages, cells were washed with phosphate buffered saline (PBS, Eurobio) and detached with Trypsin (Gibco Life Technologies #12605010) for 10 minutes at 37°C. The compound library was purchased from Prestwick Chemicals V3 and corresponds to a unique collection of 1280 off-patent small molecules, mostly approved drugs FDA, EMA and other agencies. All chemicals compounds were diluted in DMSO as 10 mM stock solution, and represent four 384-well plates. Cells were counted using T4 Cellometer (Nexcellom) and optimum cell densities were obtained as 2500 cells/well and 4000 cells/well for Nluc-CD63 and Nluc-CD9, respectively. The screening was performed at same early cell passages for both replicate experiments according to the optimized amount of cells seeded in 384-well plates (ViewPlate-384 Black Perkin Elmer, #6007460) using a Multidrop Combi (Thermo Fisher Scientific) in 40μl of total cell media.

Around 24 hours after cell seeding, cell media was removed from the plates and cells were twashed once with 40μL of phenol red free, serum free Leibovitz’s L-15 medium. A total of 40μL of the same Leibovitz’s L-15 medium was robotically added to the plates (MCA 384, Tecan). Briefly, 2μl/well of each compound at 2mM were mixed in a pre-dilution plate containing 78μl/well of cell medium, and 10μl of this solution were dispensed into each 384-cell plate well, in order to obtain a final concentration of 10μM and 0,5% of DMSO. DMSO was present in columns 1, 2, 23 and 24, and represents internal plate solvent controls. The screening was performed in two biological replicates for both cell lines.

#### Nluc assay and cell labelling

After 24 hours, 40μL of medium were transferred from cell plates to V-shaped 384-well plates using the MCA-384 head. Plates were then centrifuged at 350g for 10 min at RT. 30μL of centrifuged supernatant was then transferred to flat bottom plates with the MCA-384 head. Nanoluc reagent was freshly reconstituted according to manufacturer’s instructions (Nano-Glo® Luciferase Assay System, Promega). 5μL of reconstituted solution were added to each well using a MultiDrop Combi. Plates were shaken for 30 seconds at 300 rpm on an orbital shaker (Titramax 100, Heidolph) prior to reading. Luminescence was recorded using a CLARIOStar (BMG Labtech) (gain = 3600). In the meantime, cells were processed for nuclei labelling performed as follows: cells were fixed in a 3% formaldehyde solution for 15 min using the MCA 384 followed by 1hour incubation with the dye Hoechst 33342 (1:500, Sigma, #14533). Then PBS solution was added on top of it. Plates were kept at 4°C for 72 h prior to image acquisition.

#### Images acquisition and analysis

Image acquisition of Hoechst 33342 fluorescent nuclei (excitation: 361nm to 497nm; emission 460nm to 490nm) was performed using the INCell analyzer 6500HS automated system (GE Healthcare, USA) at a 10X magnification (Nikon 10X/0.451, Plan Apo, CFI/60), using the same exposure time for all plates in the experiment and across replicate experiments. Plates were loaded onto the microscope system with Kinedx robotic arm (PAA, UK). 16-bit images of four different positions in each well were acquired. The total number of cells measured in a field was typically around 350. For the screen, a total of 3,072 (8 × 384-well plates) wells were imaged, resulting in 12.288 grey scale images (3,072 × 4 fields of view) per replicate experiment. We then made use of morphological characteristics of Hoechst-labelled nuclei to distinguish dead cells [58] using INCell Analyzer 3.7 Workstation software (GE Healthcare). Cells were classified as alive or dead and % of dead cells computed for each drug-treated and control DMSO wells.

#### Data analysis and hit calling

Screening data quality was graphical reviewed as scatter plots and plate heat-maps to depict any bias or technical issues using in-house tools (Biophenics platform, Institut Curie). Raw luminescence signals and cell count were first log transformed before Tukey’s two-way median polishing [59] [60], then normalized as follows: sample median and median absolute deviation (MAD) were calculated from the population of each internal plate data points (named as Ref pop) and used to compute Robust Z-scores (RZ-scores, from [61]) according to the formula: *RZ*−*score*= [*compound value*−*median* (*Ref pop*)]/ [1.4826× *MAD* (*Ref pop*)] where *compound value* corresponds to the drug-treated data point, where MAD is defined as the median of the absolute deviation from the median of the tested wells. Median polished cell count values were scaled as: *Proliferation* (%)=100× [*compound value*]/[me*dian* (*negative control*)]. Hits were identified per replicate experiment as those compounds modulating EV secretion (threshold applied is: |*RZ*-*score*|>2), in two replicate experiments). Hit values were computed as median values in the final hit list. Selected compounds were subsequently annotated in regards to the Proliferation (%) and % Dead cells, in order to identify compounds which could affect cell viability.

### EV isolation and drug treatment

For EV isolation, 6*10^6 of MDA-MB-231 parental cells were seeded per 15 cm culture dish in an optimized number of dishes to obtain 70*10^6 (Fig.1), 27*10^6 (Fig. 3) or 50*10^6 (Fig.4) secreting cells. After 24 hours, cells were washed 1 X with PBS (Eurobio) and 15mL of serum free, phenol red free Leibovitz’s L-15 were added. For experiments in Fig.3 and Fig.4 DMSO (0,1%), Homosalate (10μM) or Bafilomycin A1 (100nM) were added to phenol-red free, serum free Leibovitz’s L-15 medium incubated on cells during 24 hours (DMSO and Homosalate) or 16 hours (Bafilomycin A1). The day after, cells were counted and the percentage of cell viability was determined using Trypan Blue (Invitrogen). Experiments were performed with cells showing ≥ 75% viability. In parallel, conditioned media were harvested and centrifuged at 300g for 10 min at 4 °C to remove dead cells and debris. Then, resulting supernatant was centrifuged at 2000g for 20 min at 4 °C to discard large EVs in the 2000K (=2K) pellet and then concentrated on a sterilized Sartorius Centrifugal Filter (MWCO = 10 kDa; Sartorious, #VS2001) or Centricon Plus-70 Centrifugal Filter (MWCO = 10 kDa; Millipore, #UFC701008). Media concentrated to ∼500μl were overlaid on 35 nm qEV size-exclusion columns (Izon, SP5) for separation. According to manufacturer’s instructions, SEC fractions were collected in 500μl volume. For experiments in Fig.1B and 1C, we collected SEC fractions one by one from F1 to F24. Then, not concentrated SEC fractions were individually analyzed in terms of particle number using Nanoparticle Tracking Analysis (NTA) (Particle Metrix ZetaView®), protein concentration using BCA (Pierce™ BCA Protein Assay Kit, Thermo Scientific) or Nluc activity using Nano-Glo® Luciferase Assay System (Promega). When comparing drug-treated versus DMSO negative control cells (Fig.3 and Fig.4), concentrated conditioned media from same number of secreting cells were used for SEC. In this case, SEC fractions collected in 500μL volume were pooled as F7-11 (EV-rich=2,5mL) or as F12-24 (protein-rich=6,5mL). Pooled fractions were further concentrated using 10KDa cut-off filters (Amicon Ultra-15, Millipore) before NTA, Western Blot or Transmission Electron Microscopy (TEM). To compare the effect of drug treatments on particle secretion from different tumor cell lines (MCF7, HeLa or Jurkat, Suppl.Fig.3) we adopted a procedure similar to what was described before, except that cells were seeded at a density of 2,2*10^6 (HeLa), 4,4*10^6 (MCF7) into 10 cm cell culture dishes or 10*10^6 in T25 flasks (Jurkat). Drugs were added in serum free DMEM or RPMI (Jurkat). Particle number from same number of secreting cells was then measured by NTA directly in conditioned media after 300g followed by 2000g centrifugations and concentration using sterilized Sartorius Centrifugal Filter (MWCO = 10 kDa; Sartorious, #VS2001).

### Nanoparticle tracking analysis (NTA)

NTA was performed using ZetaView PMX-120 (Particle Metrix) equipped with a 488nm laser, at x10 magnification, with software version 8.05.02. The instrument settings were 22°C, gain of 26 and shutter of 70. Measurements were done at 11 different positions (2 cycles per position) and frame rate of 30 frames per second. Image evaluation was done on particles with Minimum Brightness: 20, Minimum Area: 10, Maximum Area: 500, Maximum Brightness: 255. Tracking Radius2 was 100, Minimum Tracelength: 7.

### Western Blot

Cell lysates (CL) for Western Blot were obtained by incubating 1*10^6 cells in 25 μL of lysis buffer (50 mM Tris, pH 7.5, 0.15 M NaCl, 1% Triton X-100) with 2% complete protease inhibitor (Roche) for 15 min on ice, followed by a 13000 rpm centrifugation for 20 min at 4°C to recover the supernatant. The amount of CL used for Western Blot was the equivalent of 200 000 cells. For EVs isolated in F7-11, the amount used for Western Blot was adjusted either by same number of secreting cells (Fig. 3: 2,7*10^6; Fig.4: 5*10^6 cells) or by same number of particles (4*10^8 measured in NTA). For EVs isolated in singularly collected fractions, the used amount corresponded to 25μL of the 500μL of each not-concentrated fraction from 70*10^6 secreting cells. Samples were mixed with Laemmli sample buffer (BioRad) without β-mercapto-ethanol. After boiling 5 min at 95°C, samples were loaded on a 4-15% Mini-protean TGX-stain free gels (BioRad). Transfer was performed on Immuno-Blot PVDF membranes (BioRad), with the Trans-blot turbo transfer system (BioRad) during 7 minutes. Blocking was performed during 30 minutes with Roche blocking solution in TBS 0,1% Tween. Primary antibodies were incubated overnight at 4°C and secondary antibodies during 1h at room temperature (RT). Development was performed using Clarity western ECL substrate (BioRad) and the Chemidoc Touch imager (BioRad). Membrane were incubated with the following antibodies: mouse anti-human SLC3A2/CD98 1/3000 (clone 2B10F5 ProteinTech), mouse anti-human CD63 1/1000 (clone H5C6, BD Bioscience), mouse anti-human CD9 1/1000 (clone MM2/57, Millipore), mouse anti-human CD81 1:1000 (clone TS81, Diaclone), rabbit anti-human 14-3-3 1/1000 (EPR6380, GeneTex), rabbit anti-human Lamp1 1/1000 (clone EPR4204, GeneTex), mouse anti-Gapdh (clone 1E6D9, ProteinTech). Monoclonal rabbit anti-human Syntenin (used 1/1000) was a gift from P. Zimmermann. Secondary antibodies: HRP-conjugated goat anti-rabbit IgG (H+L) and HRP conjugated goat anti-mouse IgG (H+L) were purchased from Jackson Immuno-Research.

### Immunofluorescence

The day before the treatment, 100 000 MDA-MB-231 cells were seeded on 12 mm diameter coverslips coated with polyornithine (15 μg/mL). The cells were treated with 0,1% DMSO for 24h, 10 μM homosalate in 0,1% DMSO for 24h or 100 nM bafilomycin in 0,1% DMSO for 16h. After the treatments, they were fixed with 4% paraformaldehyde (PFA) (EMS) during 15 min at RT. The cells were incubated for 1h in a blocking solution: PBS containing 0.5% saponin and 0.1% BSA. Then primary and secondary antibodies were successively incubated during 1 h each at RT in PBS containing 0.1% saponin and 0.1% BSA. Coverslips were then mounted on slides with Fluoromount G (Invitrogen). Images were acquired on a Zeiss LSM 780 confocal microscope using an alpha Plan-Apochromat 63x/1.46 Oil with the following acquisition parameters: average per line 2, pixel size depending on the sample between 81 and 106 nm, z-step 0.33 μm for stack imaging. At least 2 fields were captured to image a total of at least 10 cells per experiment (three independent experiments) with a minimum of 54 cells per treatment. Image analysis was performed with ImageJ. A median filter of 2-pixel radius was first applied to remove noise, then a subtract background with a 20 rolling radius was done. Then, to measure colocalization between two channels, JACoP plug-in [62] was used to calculate Mander’s coefficients in each individual cell. Signal threshold Intensity to measure Mander’s coefficients was calculated with the Otsu’s method [63]. Coverlips were incubated with the following primary antibodies: SLC3A2/CD98 antibody (clone 590559 1/500, R&D Systems), mouse IgG2b anti-human CD63 (clone TS63b 1/100, available upon request to E. Rubinstein) and mouse IgG1 anti-human CD9 (clone TS9 1/100) (commercially available at Diaclone or Abcam) and then with the following secondary antibodies: goat anti-human IgG (H+L) Alexafluor 488 (Invitrogen, 1/200), goat anti-mouse IgG2b Alexafluor 647 (Invitrogen, 1/200) and goat anti-mouse IgG1 Alexafluor 568 (Invitrogen, 1/200).

### EV immunoprecipitation

For EV immunoprecipitation, the material corresponding to 1*10^9 EVs isolated in F7-11 was used for each sample. Exosome Isolation kit beads (Miltenyi) were used following manufacturer’s instructions. Briefly, EVs in F7-11 were incubated with 50μL of anti-CD63 or anti-CD9 beads overnight at 4°C. The day after, washes were performed on the columns using the isolation buffer provided in the kit. After addition of magnetically labeled EVs on columns, flow-through (FT) was recovered in the resulting running liquid which was pooled with the first wash of beads. Collected FT were then subjected to ultracentrifugation for 2 hours at 200 000g using the TLA 45 rotor (Beckman Coulter) and resuspended in 20μL of Laemmli 1X (BioRad). Elution of immunoprecipitated (IP) EVs was performed with 25μL Laemmli 1,5X. All the recovered materials of IP and FT were loaded on gels.

### Transmission electron microscopy (TEM)

Electron microscopy was performed on EVs isolated in F7-11 from same number of secreting cells (Fig. 1F-G: 7*10^6, Fig. 3C: 2,7*10^6) and stored at −80 °C that had never been thawed and re-frozen. F7-11 was deposited on formvar/carbon–coated copper/palladium grids and adsorbed for 20 min before uranyl/acetate contrasting and methyl-cellulose embedding for whole-mount analysis as described previously [64]. Staining with CD63 or CD9 antibodies was performed according to the Protein A-gold method [65] on EVs adsorbed to formvar/carbon– coated copper/palladium grids. CD63 staining was performed by incubating with mouse anti-CD63 (TS63 Diaclone 857.770.000 1/200) in PBS-BSA 1% for 30 min and CD9 staining was performed incubating with rabbit anti-CD9 (Abcam ab236630 1/80) for 30 min, 10 nm protein-A-gold (CMC, Utrecht, The Netherlands) for 20 min, fixed for 5 min with 1% glutaraldehyde (Electron Microscopy Sciences).

Subsequently, after a wash on 10 droplets of distilled water, grids were transferred to droplets of 0.4% (w/v) uranyl acetate (UA) staining and 1.8% (w/v) methylcellulose embedding solution. After 10 min of incubation, grids were picked up in a wire loop. Most of the excess of the viscous embedding solution was drained away with filter paper after which the grids were air-dried forming a thin layer of embedding solution. Images were acquired with a digital camera Quemesa (EMSIS GmbH, Münster, Germany) mounted on a Tecnai Spirit transmission electron microscope (FEI Company) operated at 80kV. EVs concentrations were estimated from digital images by counting the number of EVs per μm2. This was performed by using the ImageJ software.

### EV uptake, in vitro proliferation and in vitro anoikis resistance assay

For EV uptake, the material corresponding to 5*10^8 EVs isolated in F7-11 from MDA-MB-231 cells was used. Briefly, MCF7 recipient cells were seeded in 24 well plates at the density of 150 000 cells. After 24 hours, cells were washed 1X with PBS and serum free DMEM containing EVs was incubated on cells during 1 hour. After this time, cells were harvested by trypsinization. For proliferation assay, cells were resuspended in DMEM with 10% of FCS and transferred into xCELLigence microplates (E-plate 16, Agilent) at the density of 20 000 cells for real-time analysis of cell adhesion and growth. Plates were loaded into xCELLingence RTCA DP instrument (Agilent) inside a 37°C incubator. A run of 50 hours with readings every 30 minutes was programmed and the slopes in the range 10-40h were calculated using Roche RTA software. For the in vitro anoikis assay, cells collected after trypsinization were resuspended in serum free DMEM containing 0,1% BSA and kept on ultra-low attachment (ULA) six-well plates (Corning, #3471) for 24 h. Next, cells were collected and washed with PBS, treated for 5 min with trypsin for the disruption of cell aggregates and transferred into xCELLigence microplates following the same procedure described for proliferation assay.

### Statistical analysis

Following the recommendation of D.L.Vaux [66], for each experiment where number of biological replicates were 2 or 3, we displayed the results in a transparent manner, showing each individual biological replicate as a dot, so the readers could interpret the data for themselves. We (as suggested by D.L. Vaux) considered that same trends of results obtained independently 2-3 times were as informative as statistical tests to evaluate reproducibility of the experiments. Nonetheless, we also performed statistical analyses with GraphPad Prism version 8.0.2 (GraphPad software, California USA), by paired t-test (Fig.3B; Fig. 4B; Suppl. Fig. 3B), Mann-Whitney test (Fig. 3C), Ordinary one way Anova (Fig. 4E, Fig. 5A and 5B).

## Acknowledgments

We thank for fruitful discussions several team members, especially Dr Mercedes Tkach Mathilde Mathieu, and Emeline Bonsergent, Federico Cocozza, Davinia Arguedas and Alix Zhou, and for helpful discussions and tools Drs Michael Boutros and Alena Ivanova (DKFZ, Germany), Dr Michael Davidson (The Florida State University, USA), Dr Walter Mothes (Yale University School of Medicine, USA), Dr Eric Rubinstein (Centre d’Immunologie et des Maladies Infectieuses, France) and Dr. Pascale Zimmermann (Centre de Recherche en Cancerologie de Marseille, France and KU Leuven, Leuven, Belgium). We also acknowledge the following platforms of Institut Curie: PICT-IBiSA, member of the France-BioImaging national research infrastructure (ANR-10-INBS-04) for fluorescence and electron microscopy, and the genomic platform for the authentication of cell lines by STR. This work was funded by INSERM, CNRS, Institut Curie, French IdEx and LabEx (ANR-10-IDEX-0001-02 PSL, ANR-10-LABX-0038, ANR-11-LABX-0043, ANR-18-IDEX-0001 Université de Paris), grants from french ANR (ANR-18-CE13-0017-03; ANR-18-CE15-0008-01; ANR-18-CE16-0022-02), INCa (INCA-11548), Fondation ARC (PGA1 RF20180206962), FRM (SPF20170938694), USA NIDA (DA040385). The BioPhenics laboratory is supported by Institut Curie, IBiSA (PICT-IBiSA) and part of ChemBioFRance and France-BioImaging national infrastructures.

## Authors contribution

E.G, L.MJ. and C.T. designed the study, interpreted the data, wrote the article. E.D.N contributed to screening design and implementation. E.G, N.N., A.L., M.J. performed experiments. M.C. designed and implemented the computational framework to analyse the screening data. E.G, N.N., A.L., M.C., M.J. analyzed the data. E.G, A.J., G.L. generated plasmids or cells. A.J. and E.D.N interpreted data. All authors read and corrected the article.

---

**Supplementary Figure 1.**
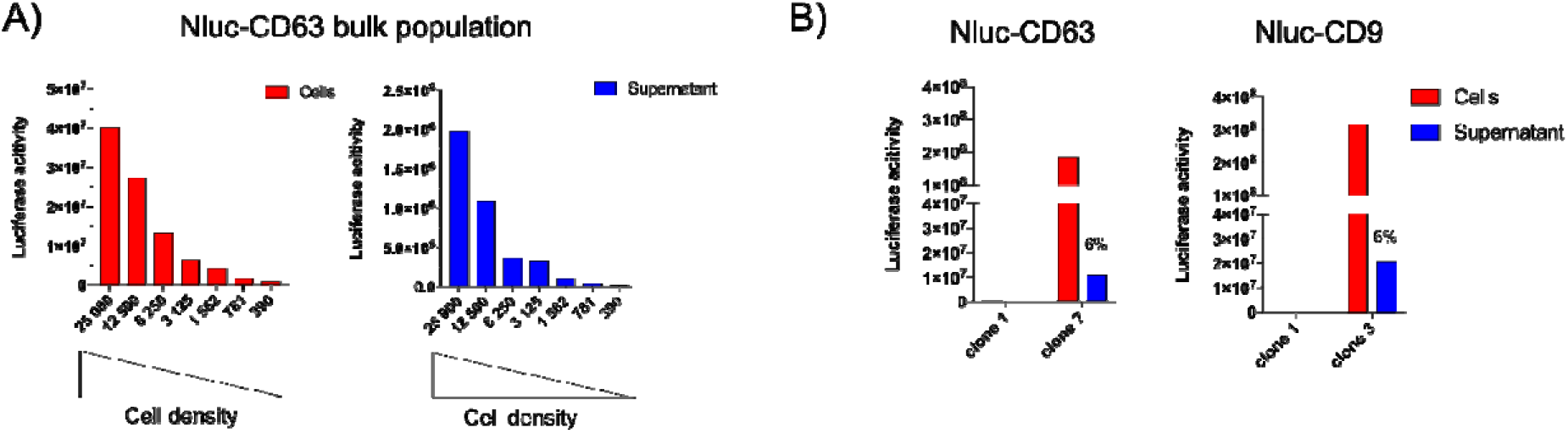
Nanoluciferase tagged CD63 and CD9 are secreted into EVs. A) Dose-scale experiment to determine the limit of detectability of Nluc activity in cells (red) or supernatants (blue) from Nluc-CD63 stable transfected bulk population. Serial 1:2 dilutions of cells starting from 25000 cells were seeded in 96 well plates and Nluc activity in cells or supernatants was measured 24 hours later. Shown data are from a single pilot experiment. B) Selection of stable clonal populations transfected with Nluc-CD63 (left panel) or Nluc-CD9 (right panel) by measurement of Nluc-activity in the cells (red) or in the supernatant (blue). Measurements of Nluc activity in cells and supernatants for each clone were done from 25000-30000 cells. Shown percentages represent the ratios between supernatant versus cell Nluc activity.

**Supplementary Figure 2.**
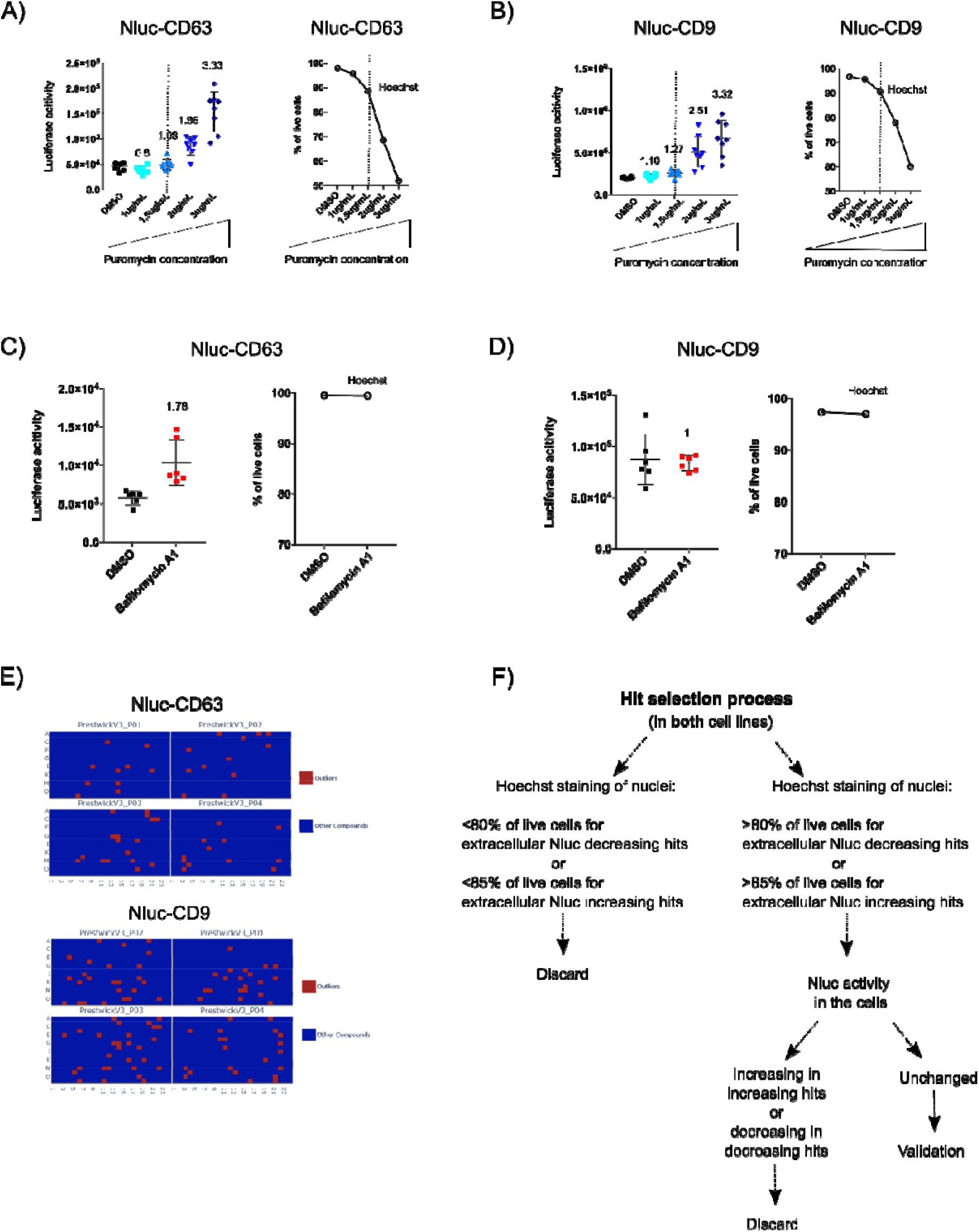
Identification of a drug increasing extracellular Nluc activity in Nluc-CD63 and Nluc-CD9 cells. A) and B) Dose-scale experiment to determine the effect of Puromycin-related toxicity on the measurement of extracellular Nluc activity in Nluc-CD63 or Nluc-CD9 cells. Increasing doses of Puromycin (1μg/mL; 1.5μg/mL; 2μg/mL;3μg/mL) were administered to 2500 Nluc-CD63 cells (A) or 4000 Nluc-CD9 cells (B). For both cell lines, extracellular Nluc activity was measured and reported as ratio on DMSO negative control. Hoechst staining of nuclei was used on the same cells to determine the percentage of live cells. Data are from a single pilot experiment. C) and D) Determination of Bafilomycin A1 effect on the measurement of extracellular Nluc activity in Nluc-CD63 and Nluc-CD9 cells. 2500 Nluc-CD63 cells (C) or 4000 Nluc-CD9 cells (D) were treated with 100nM Bafilomycin A1 for 16h. For both cell lines, extracellular Nluc activity was measured and reported as ratio on DMSO negative control. Hoechst staining of nuclei was used on the same cells to determine the percentage of live cells after Bafilomycin A1 treatment. Data are from a single pilot experiment. E) Schematic representation of screening results for Nluc-CD63 (above) or Nluc-CD9 (below). For each cell line, the four rectangles done of blue spots represent the four 384 well plates composing the entire drug library. Red spots represent outlier compounds which affected Nluc activity in the supernatant compared to DMSO negative control. Intensity of Nluc activity is expressed as robust Z-score= [(compound value-median of (Ref pop))/ (MADnc X 1.4826)], MAD= [median (|Ref pop-median (Ref pop)|)]. Increasing or decreasing hits were called according to the Threshold: |Robust Z score|>2 or <-2. Data from two independent experiments are shown. F) Scheme of hit selection process.

**Supplementary Figure 3.**
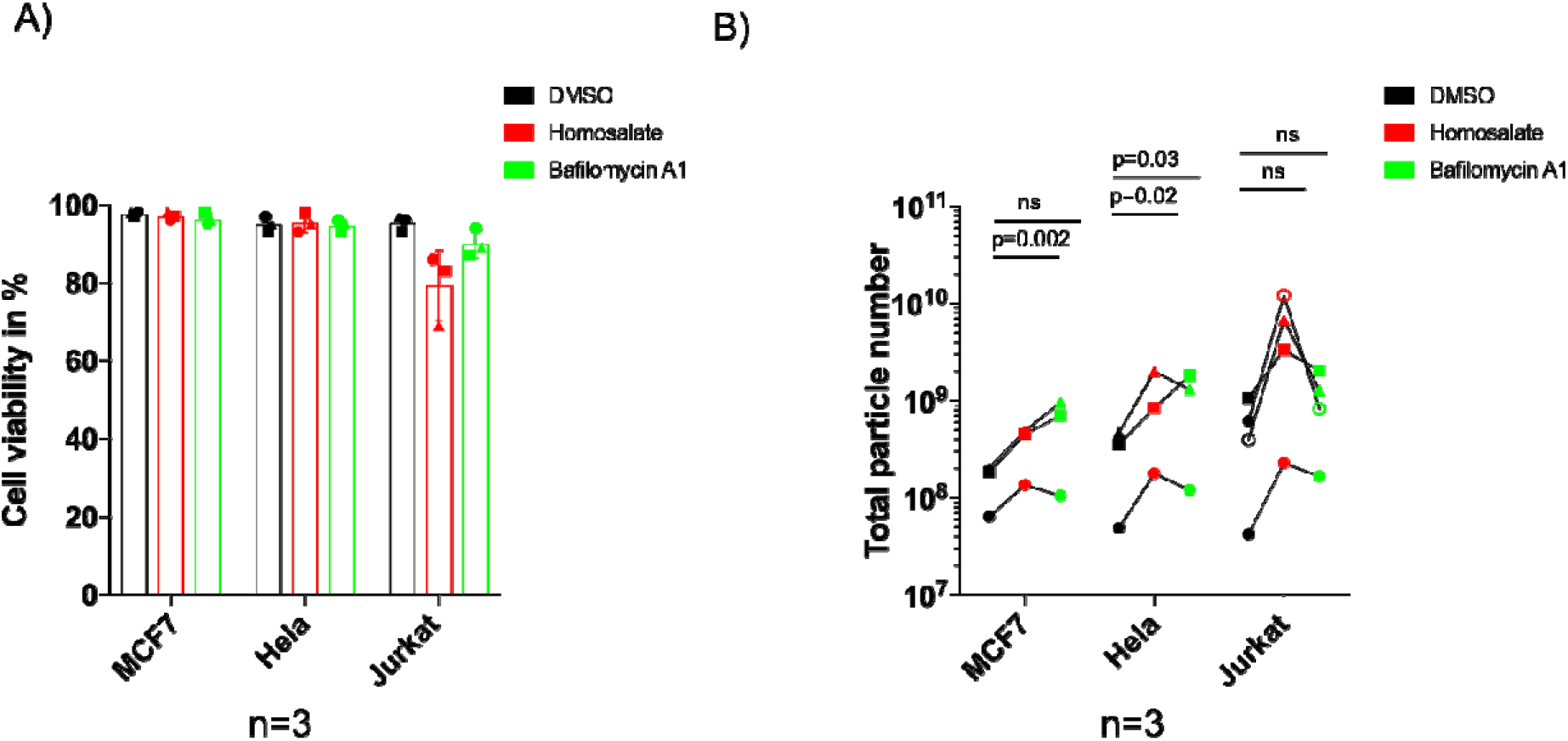
Homosalate increases particle secretion in tumor cell lines. A) Quantification of cell viability after Homosalate and Bafilomycin A1 treatment in MCF7, Hela and Jurkat cells. DMSO, Homosalate or Bafilomycin A1 treated cells were counted after conditioned media collection using Trypan Blue as a reporter of cell death. Cell viability is expressed in percentage. Jurkat cells reproducibly displayed less than 85% viability upon Homosalate treatment. Data from three or four (Jurkat) independent experiments are shown Quantification of particles induced by treatment with Homosalate or Bafilomycin A1 in MCF7, Hela and Jurkat cells. Graphs show total particle number recovered in concentrated conditioned media from 3*10^6 cells, measured by NTA. Data from three or four (Jurkat) independent experiments are shown. Paired parametric t-test: MCF7 p=0.002, ns; Hela p=0.02, p=0.03, Jurkat ns.

